# Impact of adolescent intermittent ethanol exposure on interneurons and their surrounding perineuronal nets in adulthood

**DOI:** 10.1101/2022.01.07.475220

**Authors:** Carol A. Dannenhoffer, Alexander Gómez-A, Victoria A. Macht, Rayyanoor Jawad, E. Blake Sutherland, Ryan P. Vetreno, Fulton T. Crews, Charlotte A. Boettiger, Donita L. Robinson

## Abstract

**Background:** Binge alcohol exposure during adolescence results in long-lasting alterations in brain and behavior. For example, adolescent intermittent ethanol (AIE) exposure in rodents results in long-term loss of functional connectivity among prefrontal cortex (PFC) and striatal regions as well as a variety of neurochemical, molecular, and epigenetic alterations. Interneurons in the PFC and striatum play critical roles in behavioral flexibility and functional connectivity. For example, parvalbumin (PV) interneurons are known to contribute to neural synchrony, and cholinergic interneurons contribute to strategy selection. Furthermore, extracellular perineuronal nets (PNNs) surround some interneurons, particularly PV+ interneurons, to further regulate cellular plasticity. The effect of AIE exposure on expression of these markers within the PFC is not well understood.

**Methods:** The present study tested the hypothesis that AIE exposure reduces expression of PV+ and ChAT+ interneurons in the adult PFC and striatum and increases related expression of PNNs (marked by binding of *Wisteria Floribunda* agglutinin lectin; WFA) in adulthood. Male rats were exposed to AIE (5 g/kg/day, 2-days-on/2-days-off, *i*.*g*., P25-P54) or water (CON), and brain tissue was harvested in adulthood (> P80). Immunohistochemistry and co-immunofluorescence were used to assess expression of ChAT, PV, and WFA labeling within the adult PFC and striatum following AIE exposure.

**Results:** ChAT and PV interneuron numbers in the striatum and PFC were unchanged after AIE exposure. However, WFA labeling in the PFC of AIE-exposed rats was increased compared to CON rats. Moreover, significantly more PV neurons were surrounded by WFA labeling in AIE-exposed subjects relative to controls in both PFC subregions assessed: the orbitofrontal cortex (CON = 34%; AIE = 40%) and the medial PFC (CON = 10%; AIE = 14%).

**Conclusions:** These findings indicate that while PV interneuron expression in the adult PFC and striatum is unaltered following AIE exposure, PNNs surrounding these neurons (indicated by extracellular WFA binding) are increased. This increase in PNNs may restrict plasticity of the ensheathed neurons, thus contributing to impaired microcircuitry in frontostriatal connectivity and related behavioral impairments.

## Introduction

In humans, alcohol use is commonly initiated during adolescence (Spear, 2000), with over 7 million individuals aged 12 to 20 having reported drinking in the past month and over half of these reporting binge drinking (4-5 drinks in a 2-hour period; SAMHSA, 2020). As the brain undergoes critical neurodevelopment during adolescence, initiation of alcohol use during this time has long-term consequences on neural function and behavior (Crews et al., 2019; Spear, 2000; Spear, 2016; Spear, 2018; Spear & Swartzwelder, 2014). In fact, both human (Scaife & Duka, 2009; Yoo & Kim, 2016) and rodent (Coleman Jr et al., 2014; Fernandez et al., 2017; Varlinskaya et al., 2020) studies have found that adolescent binge alcohol exposure is detrimental to performance in adulthood on behavioral tasks requiring cognitive flexibility (Gass et al., 2014; Macht et al., 2020; Sey et al., 2019 and see Crews et al., 2019; Crews et al., 2016 for reviews).

Behavioral flexibility is regulated by a complex circuit with the prefrontal cortex (PFC) exerting top-down control over subcortical areas (see Dannenhoffer et al., 2021 for review). Human studies show that individuals who abuse alcohol exhibit reduced frontostriatal connectivity, particularly between the dorsolateral PFC and limbic portion of the striatum (Elton et al., 2021). Rodent models have highlighted that this circuit is disrupted such that cognitive flexibility deficits following adolescent intermittent ethanol (AIE) exposure are associated with a loss of frontostriatal connectivity (Gómez-A et al., 2021). However, the cellular mechanisms underlying this loss of frontostriatal connectivity are poorly understood.

Cholinergic neurons, which express choline acetyltransferase (ChAT) along with other markers, are particularly sensitive to developmental influence of alcohol. ChAT-positive neurons in the basal forebrain project to the PFC and, thus, are key integrators of cortico-subcortical connectivity (Mayo et al., 1984; Mesulam et al., 1983). Studies have found that AIE exposure reduces ChAT expression in both striatal interneurons (Galaj et al., 2019; Vetreno et al., 2014) and basal forebrain projection neurons (Coleman Jr et al., 2011; Vetreno et al., 2014). Parvalbumin (PV)-positive interneurons are also sensitive to the effects of alcohol during adolescence. PV interneurons release GABA and provide critical inhibitory regulatory control during behavioral flexibility tasks such that inactivation of these neurons disrupts behavior in working memory and cognitive flexibility related tasks (Murray et al., 2015). AIE exposure has been shown to decrease PV-positive interneurons in the hippocampus (Liu & Crews, 2017), but not in the basal forebrain (Coleman Jr et al., 2011); however, it is unclear if a similar reduction in expression is observed in the PFC that may contribute to reduced frontostriatal connectivity.

AIE-induced changes in interneuron regulation of frontal and striatal circuitry may be further compounded by AIE-induced changes in surrounding extracellular matrix proteins, specifically perineuronal nets (PNNs). PNNs influence these circuits by providing stability and protection to neurons and synapses (Lasek, 2016; Reichelt et al., 2019). These nets are lattice-like structures that surround neurons and synapses, particularly PV interneurons (Baker et al., 2017; Härtig & Brauer, 1992), and regulate the intrinsic excitability of cortical interneurons (Chu et al., 2018), thus contributing to overall excitatory/inhibitory balance (Hensch, 2005). Importantly, low expression of PNNs within the PFC allow for heightened plasticity during adolescence (Drzewiecki et al., 2020) as this is a crucial PFC maturation phase (Spear, 2000). Alcohol during this time is detrimental to PNN integrity, as Coleman Jr et al. (2014) found increases in WFA-labeled PNN density in the orbitofrontal cortex (OFC) of AIE-exposed mice relative to control mice.

The current study expanded upon these previous findings by testing the hypothesis that AIE exposure disrupts ChAT and PV expression within the PFC and striatum, as well as binding of the PNN marker WFA in the surrounding extracellular space. We also tested whether AIE exposure changed the proportion of PV interneurons that were encapsulated by PNNs within subregions of the PFC in adulthood. These experiments may provide insights into AIE-induced impairments in brain regions required for executive function reliant upon frontostriatal connections.

## Methods

### Subjects

A total of 20 male Sprague-Dawley rats were used in the present experiment (CON n=9; AIE n=11). Archived tissue from a previous study (N=16; CON n=8; AIE n=8) was used for follow-up analyses; that study also used male Sprague-Dawley rats that underwent identical alcohol and water exposures. Males were used due to availability (female littermates were used in another ongoing study) and to replicate and extend previous findings following AIE exposure that were observed in adult male rodents (Coleman Jr et al., 2014; Vetreno et al., 2020). All rats were bred and reared in-house and maintained in a temperature and humidity-controlled vivarium with a 12hr:12hr light cycle (lights on at 0700) and provided *ad libitum* access to food and water. Litters were culled to 10 pups by postnatal day (P) 3. Animals were housed two per cage at time of weaning (P21). All experimental procedures were approved by the Institutional Animal Care and Use Committee of the University of North Carolina at Chapel Hill and conducted in accordance with National Institute of Health regulations for the care and use of animals in research.

### Adolescent Intermittent Ethanol (AIE) Exposure

Animals were given doses of alcohol intragastrically (*i*.*g*.; 5 g/kg of 25% v/v ethanol in water) or water at equivalent volumes (control) during adolescence (postnatal day (P) 25-54) in a two-days-on/two-days-off regimen for a total of 16 doses. These doses per body weight produce blood alcohol levels of over 230 mg/dL (Madayag et al., 2017).

### Tissue Collection and Preparation

Adult animals (P80-P85) were anesthetized with a lethal dose of urethane and transcardially perfused with 0.1 M phosphate-buffered saline (PBS) followed by 4% paraformaldehyde in PBS. Brains were extracted and post-fixed in 4% paraformaldehyde for 24 hours at 4°C and then stored in 30% sucrose in PBS until sliced. Coronal sections of 40 µm were cut on a sliding microtome (HM 450, ThermoScientific, Austin, TX), and sections were sequentially collected into 12-well plates and stored at -20°C in cryoprotectant solution (30% glycerol, 30% ethylene glycol in PBS).

### Immunohistochemistry

#### Choline Acetyltransferase (ChAT), Parvalbumin (PV), *Wisteria floribunda* agglutinin (WFA)

Every twelfth section of the regions of interest (PFC: from Bregma +5.16 mm to +2.76 mm; striatum: from Bregma +2.28 mm to -0.24 mm; basal forebrain: from Bregma +1.20 mm to +0.60 mm) was stained to capture at least 4-6 representative sections per subject, allowing for systematic random sampling across the regions from anterior to posterior. Free-floating serial sections of rat brains including PFC, striatum, and basal forebrain (Figure 1) were washed 5 × 5 min in PBS, then quenched in 0.6% H_2_O_2_ to block endogenous peroxidase activity (30 minutes), and then washed 3 × 5 min in PBS again at room temperature. Next, samples were incubated in blocking solution (<u>ChAT</u>: 4% normal rabbit serum with 0.2% Triton-X-100 in 0.1M PBS; <u>PV</u>: 5% normal goat serum with 0.2% Triton-X-100 in 0.1M PBS; <u>WFA</u>: no block). Slices were then incubated in primary antibody for 48 hours at 4°C (<u>ChAT (1:200)</u>: goat anti-ChAT (Abcam Cat. #ab144P, Millipore, Temecula, CA), <u>PV (1:2000)</u>: rabbit anti-PV (Abcam Cat. #ab11427, Pittsburgh, PA); <u>WFA (1:100)</u>: WFA lectin (Sigma Aldrich Cat. #L1516, St. Louis, MO)), and then diluted in 5% normal serum with 0.2% Triton-X-100 in 0.1M PBS. Sections were then washed 3 × 5 min with PBS, incubated in appropriate secondary antibody solutions for 2 hours (<u>ChAT (1:200)</u>: rabbit anti-goat IgG (biotinylated; Vector Laboratories, BA-5000); <u>PV (1:200)</u>: goat anti-rabbit IgG (unconjugated; AB-2337913 Jackson ImmunoResearch Laboratories Inc., West Grove, PA); <u>WFA</u>: no secondary). Slices were then incubated in avidin-biotin complex (ABC; Vectastain® ABC Kit; Vector Laboratories) solution (ThermoFisher Scientific, Cat. #32020) at room temperature for one hour, as directed by the manufacturer instructions. Immunoreactivity was visualized by staining sections with 3,3’-diaminobenzidine tetrahydrochloride (DAB), which was activated by H_2_O_2_. Slices were then mounted, dehydrated, and cover slipped.

**Figure 1.**
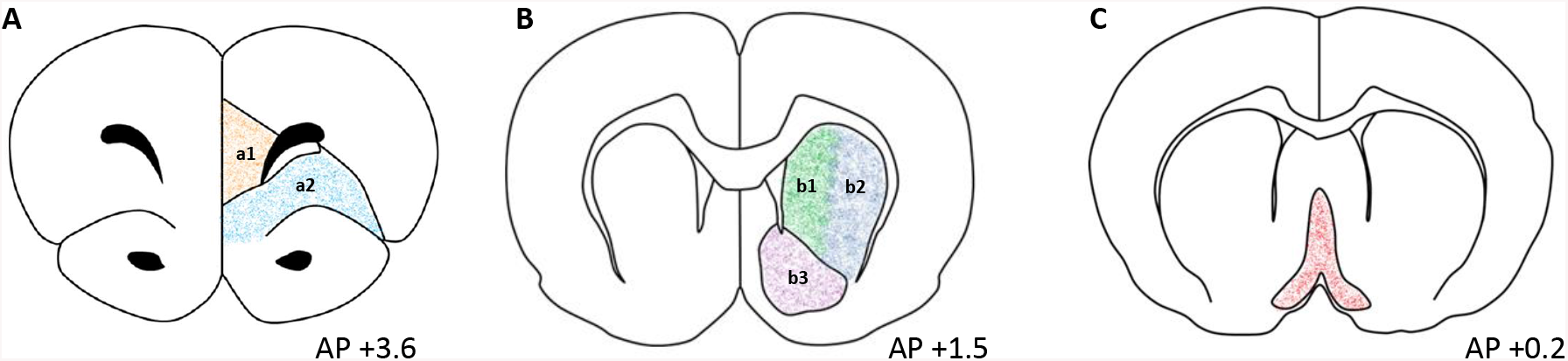
Schematic of targeted brain regions. Representative graphs of targeted brain regions including (A) orbitofrontal cortex (OFC; orange) and medial prefrontal cortex (mPFC (a1); teal); (B) dorsomedial striatum (DMS; green; b1), dorsolateral striatum (DLS; blue; b2), ventral striatum (VS; purple; b3); and (C) basal forebrain (BF; red). Coordinates are relative to bregma as defined by Paxinos and Watson (2006). AP = anterior posterior.

### Immunofluorescence

#### Parvalbumin (PV) and *Wisteria floribunda* agglutinin (WFA)

As PNNs preferentially ensheathe PV interneurons in cortical areas of the brain (Baker et al., 2017), we performed co-label immunofluorescence to determine the proportion of WFA-wrapped PV interneurons within the mPFC (Fig. 1A a1) and OFC (Fig. 1A a2); we did not assess WFA in the striatum or basal forebrain as little WFA+IR was detected in these regions (see Supplemental Figures 5 (striatum) and 6 (basal forebrain)). Every twelfth section from Bregma +5.16 mm to +2.76 mm (according to Paxinos and Watson (2006)) were stained to capture at least 4-6 representative sections per subject allowing for systematic random sampling across the PFC from anterior to posterior. Slices were washed 3 × 5 min in PBS, and then placed into 5% blocking solution (5% normal Goat serum with 0.2% Triton-X-100 in 0.1M PBS) for one hour. Next, slices were incubated in a primary antibody blocking solution including both rabbit anti-PV (1:500; Abcam Cat. #ab11427, Pittsburgh, PA), and WFA lectin fluorescein (1:100; FL-1351-2, Vector Laboratories, Burlingame, CA) at 4°C for 48 hours. Following primary antibody incubation, tissue slices were washed with PBS, and incubated in secondary antibody (goat anti-rabbit Alexa Fluor 594 ((1:1000) ThermoFisher Scientific Cat. #A32740, Rockford, IL)) for two hours and washed again. Slices were then mounted, and coverslipped using ProLong™ Gold Antifade Mountant (Cat. #P36930, ThermoFisher). As preliminary staining indicated that there was no observable co-expression between ChAT and WFA, we did not assess ChAT-WFA co-labeling.

### Microscopic Quantification and Image Analysis

Immunohistochemical DAB-stained slices were visualized, and then photomicrographs of representative sections from each region were quantified by a blind observer using an Olympus BX50 Microscope at 10×1 magnification and BioQuant Nova Advanced Image Analysis System™ (R&M Biometric, Nashville, TN) software (as described in Vetreno et al., 2020 and Vetreno & Crews, 2018). A minimum of 4 sections were quantified per rat using a modified version of non-biased stereology, which we have previously validated and published (see Nixon & Crews, 2002). ChAT and PV were quantified as the total number of positive immunoreactive (+IR) cells per area. WFA was quantified as both the total number of positively immunoreactive pixels per area and total number of WFA-labeled matrices surrounding individual cell soma. For all data collection, tissue were analyzed under identical conditions to control for background noise and to avoid non-systematic variations (Vetreno et al., 2014).

A minimum of four immunofluorescent sections were visualized and quantified using a DS-RiZ scope under identical conditions (Nikon, Inc., Melville, NY) and NIS-Elements AR46 (Nikon, Inc., Melville, NY) as previously described (Vetreno et al., 2020). PV and WFA were quantified as the total number of objects (cells and matrices) per area. As WFA is an extracellular structure and PV is a calcium-binding protein primarily localized to the cell soma, co-labeled immunofluorescence was defined as PV+ cells which were surrounded by WFA labeling. Images were taken through a z-plane within the center of the tissue section, containing 9 stacks (2.5 µm/stack) from the mPFC and OFC of all subjects. Since PNNs surround neurons, collection of multiple stacks can provide a clearer and more complete image of a net around a neuron (Slaker et al., 2016). All sections were quantified by an experimenter who was blinded to the group conditions using a modified version of unbiased stereological quantification methods, as previously published (Nixon & Crews, 2002).

WFA binding was assessed using both DAB and fluorescent methods; however, it should be noted that data quantification processes differed across techniques. For example, determining the number of objects within a region was consistent between subjects using a threshold value; however, computer detection of objects was assessed differently on the Nikon microscope versus BioQuant microscope.

### Statistical Analyses

All statistical analyses were conducted using GraphPad Software 8.0 (San Diego, California). In DAB-stained tissue, dependent variables included the quantity of ChAT and PV interneurons within the basal forebrain as well as within subregions of the PFC (OFC and mPFC) and striatum [dorsomedial (DMS), dorsolateral (DLS), and ventral (VS); **Figure 1**] and were reported as cells/mm^2^. WFA binding was quantified in a similar manner (objects/mm^2^); however, pixel density (pixels/mm^2^) was also measured. Two-tailed *t*-tests were used to assess differences in expression of targeted markers in AIE-exposed and water-exposed males. Standard α levels (α = 0.05) were used to assess group differences; however, thresholds were corrected for multiple comparisons within subregions (α = 0.05/number of t-tests; striatal subregions (α = 0.0167) and prefrontal subregions (α = 0.025)).

Using immunofluorescence, we were able to visualize both PV and WFA labeling within the same tissue, allowing us to characterize the proportions of PV that were surrounded by nets and PNNs that contained PV neurons, given an intersection of these targets. We also compared the quantity of neurons (PV) and matrices (WFA) within the area (OFC, mPFC). The fluorescent data were limited to the PFC areas, as no WFA binding was reliably observed within basal forebrain or striatal sections. PV, WFA, and co-label of PV and WFA were assessed using MANOVAs within each subregion. Due to violation of Levene’s test (PV+WFA/PV α = 0.028), data in the OFC were transformed (reciprocal).

## Results

### Choline Acetyltransferase (ChAT) Immunoreactivity

We hypothesized that ChAT interneuron expression within the PFC and striatum would be reduced as others have shown striatal reductions in ChAT expression (Galaj et al., 2019; Vetreno et al., 2014). However, we found no differences between AIE-exposed rats and water-exposed controls in the striatum (*p*=0.799; **Figure 2A**) or PFC (*p*=0.422; **Figure 2B**). The same data divided into subsections (**Supplemental Figure 1**) confirmed no group differences in subregions of striatum (top panel) or PFC (bottom panel) wherein α levels were adjusted to correct for multiple comparisons. As a positive control, we replicated the loss of basal forebrain ChAT expression (*t* (17) =5.03; *p*<0.001; **Figure 2C**; Vetreno et al., 2020; Vetreno et al., 2014; Vetreno & Crews, 2018). As further confirmation, we assessed ChAT expression in the ventral striatum in a follow-up study using archived tissue and again observed no difference between CON and AIE subjects (*p*=0.255; see **Supplemental Figure 2A**). Thus, of the regions presently assessed, the AIE-induced reductions in ChAT expression were localized to the basal forebrain.

**Figure 2.**
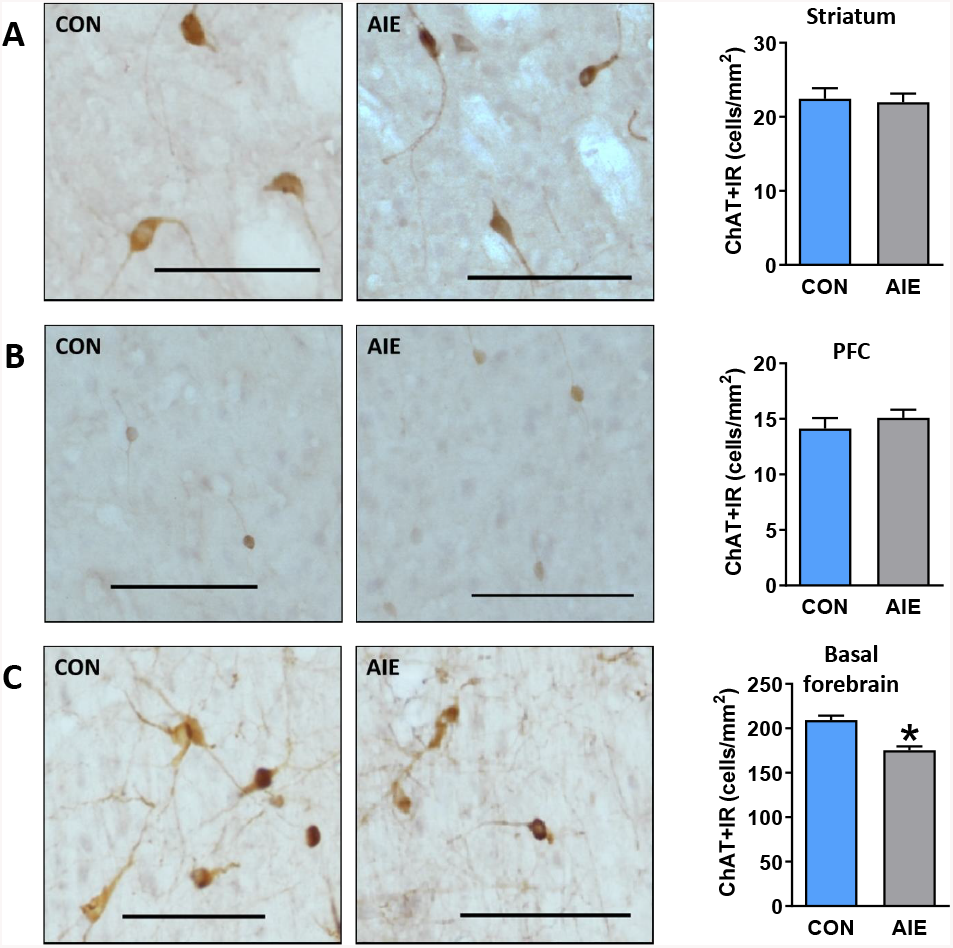
AIE-exposed rats exhibit reduced ChAT+IR in the basal forebrain, but not prefrontal or striatal regions. Representative micrographs showing immunohistochemical staining of ChAT+ cells in the (A) striatum (sample image taken from dorsolateral (DLS)), (B) prefrontal cortex (PFC; sample image taken from prelimbic (PrL)), and (C) basal forebrain of AIE and CON adult rats. We did not find reduced ChAT expression within striatal or prefrontal regions; however, as a positive control, we observed reduced ChAT expression in basal forebrain. Data are presented as mean ± SEM. * indicates main effect of exposure. Scale bar represents 100 µm.

### Parvalbumin (PV) Immunoreactivity

We next investigated PV interneurons within the same regions. We did not find differences in expression of PV+IR between the AIE-exposed and control-exposed groups in the striatum (*p*=0.170; **Figure 3A**), PFC (*p*=0.543; **Figure 3B**), or basal forebrain (*p*=0.425; Figure 3C). **Supplemental Figure 3** shows the same data separated into subregions (and adjusted α levels to correct for multiple comparisons) of the striatum (top panel) and PFC (bottom panel), where we did not observe differences between groups. Thus, AIE exposure did not alter the number of PV-positive neurons within striatum, PFC or basal forebrain slices.

**Figure 3.**
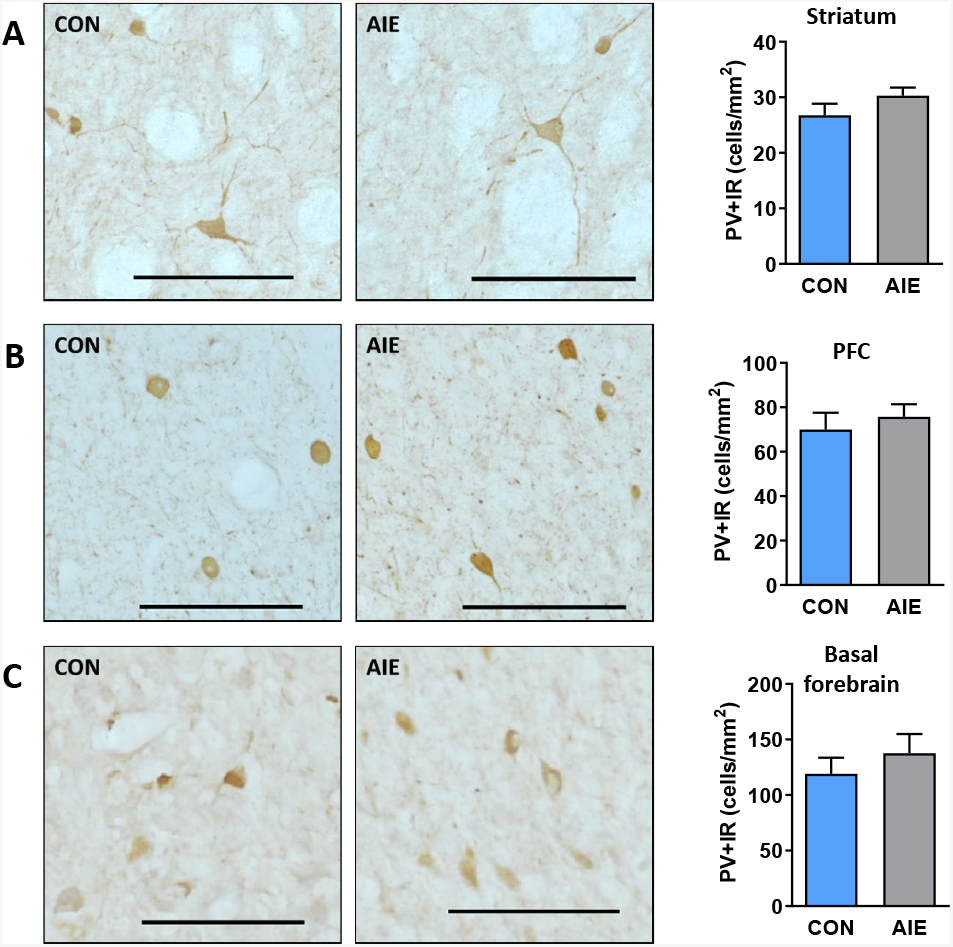
AIE exposure did not alter expression of PV+IR interneurons in the PFC, striatum, or basal forebrain of adult rats. Representative micrographs showing immunohistochemical staining of PV+ cells in the (A) striatum (sample image taken from DLS), (B) PFC (sample image taken from PrL), and (C) basal forebrain of AIE and CON adult rats. We did not find altered PV expression within any of the observed areas. Data are presented as mean ± SEM. * indicates main effect of exposure. Scale bar represents 100 µm.

### Perineuronal Net (PNN) Binding Expression

We next tested whether PNNs in the PFC were affected by AIE exposure, which could impact the function of the PV interneurons they surround. We found a significant increase in WFA labeling in AIE-exposed rats relative to control rats in both the mPFC (*t* (17) =2.56; *p*=0.02)) and OFC (*t* (17) =2.53; *p*=0.02; **Figure 4A and 4B**, respectively; α threshold = 0.025 to correct for multiple comparisons). We further subdivided the mPFC into prelimbic (PrL) and infralimbic (IL) and found a significant increase in WFA binding within the IL (*t* (17) =2.63; *p*=0.02) and a marginal increase within the PrL (*p*=0.0255; see **Supplemental Figure 4**).

**Figure 4.**
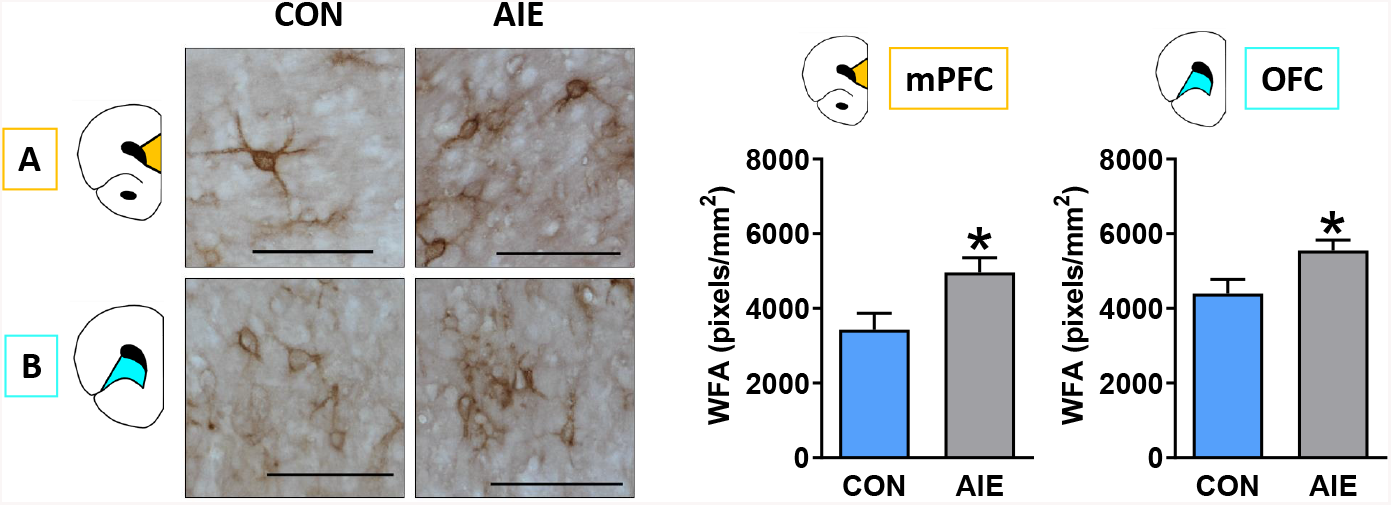
AIE exposure increased PNN density within the PFC of adult rats. Left: Representative micrographs showing immunohistochemical staining of WFA binding within the (A) mPFC and (B) OFC of CON and AIE rats. Right: WFA binding expression was significantly higher in AIE rats relative to CON in both subregions. Data are presented as mean ± SEM. * indicates main effect of exposure. Scale bar represents 100 µm.

We observed little to no WFA labeling within the striatum (see **Supplemental Figure 5**) and basal forebrain (see **Supplemental Figure 6**) in both AIE- or CON-exposed rats. Therefore, WFA data from these regions were not analyzed.

### Immunofluorescence Quantification

We hypothesized that irrespective of AIE exposure effects on PV interneuron expression, AIE exposure may functionally alter PV interneurons via modifications in extracellular PNNs, which can be detected as altered WFA binding expression. Thus, we used immunofluorescence to label both PV interneurons (in red) and WFA lectin (in green) to determine the spatial relationship between the two. To quantify PV interneuron and PNN co-expression, we calculated percentages of PV interneurons surrounded by PNNs out of total PV interneurons within the region. A MANOVA was used to compare AIE and CON subjects for three measures: PV expression, WFA binding, and percentage of WFA-enwrapped PV neurons ((PV + PNN interaction / total PV objects)*100).

Within the mPFC (**Figure 5A**), the multivariate analysis revealed a significant effect of AIE exposure to increase WFA binding expression (F (1) =11.86; *p*=0.003) and increase the percentage of WFA-enveloped PV interneurons (F (1) =10.97; *p*=0.004), but AIE exposure did not alter PV expression (*p*=0.11). Within the OFC (**Figure 5B**), the MANOVA initially violated Levene’s test (*p*=0.028); therefore, data were transformed (reciprocal) to account for group variances. The following MANOVA yielded similar results to the mPFC analysis: no effect of AIE exposure on PV expression (*p*=0.523), but significant AIE-induced increases in WFA binding expression (F (1) =7.727; *p*=0.013), and in percentage of WFA-enveloped PV interneurons (F (1) =8.240; *p*=0.011).

**Figure 5.**
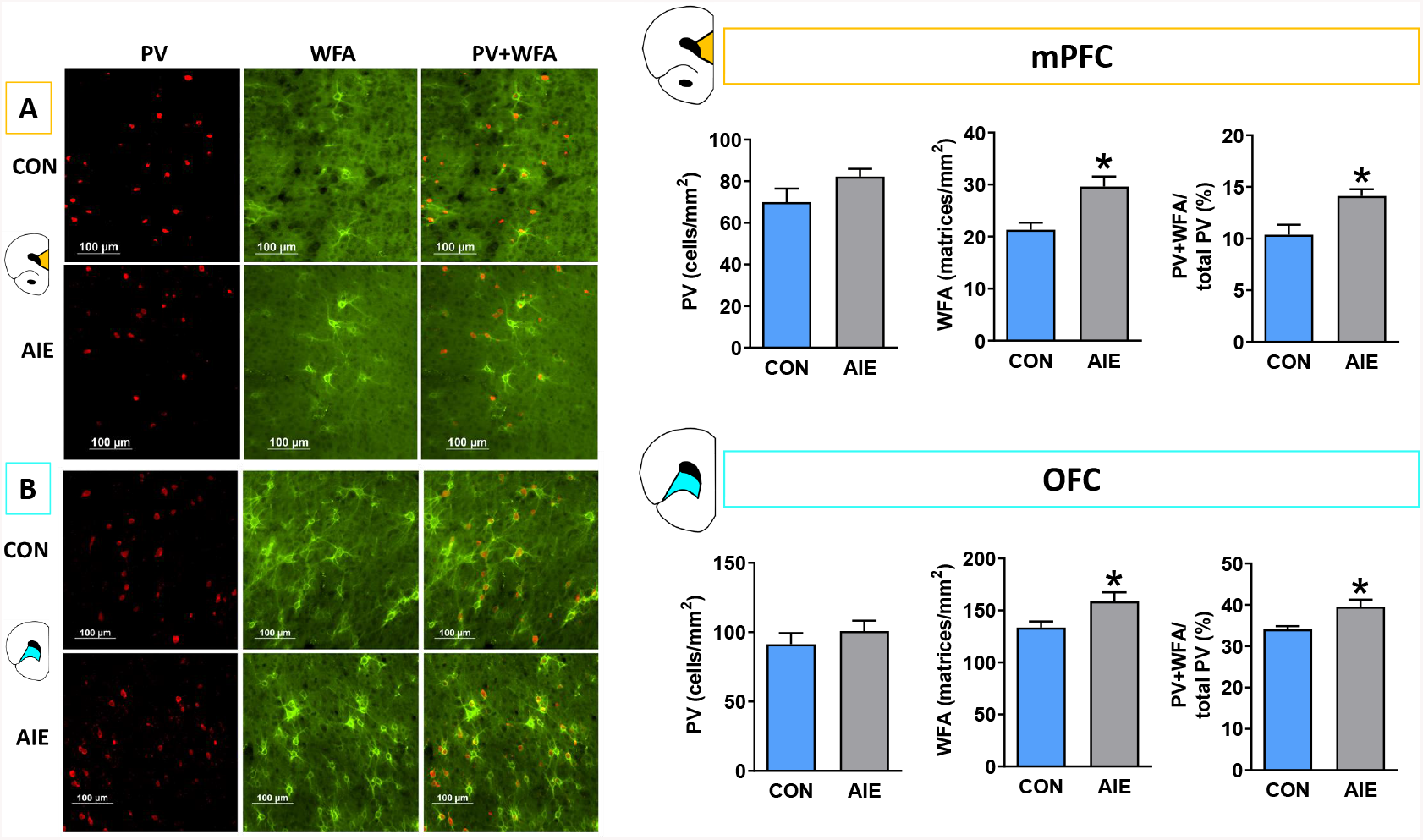
AIE-exposed rats have significantly more PNNs than controls in the PFC, and this increase is associated with PV interneurons. Left: Representative micrographs showing immunofluorescence of PV and WFA labeling within the (A) mPFC and (B) OFC of CON and AIE rats. Right: We observed no changes in PV expression in either subregion; however, WFA expression was significantly higher in AIE rats relative to CON in both subregions. In addition, a higher proportion of PV interneurons were surrounded by WFA expression in both sub regions. Data are presented as mean ± SEM. * indicates main effect of exposure. Scale bar represents 100 µm.

## Discussion

Our data show that adolescent ethanol exposure produces long-lasting increases in extracellular binding of the PNN marker WFA surrounding PV interneurons in the medial and orbital PFC, as well as reduced expression of cholinergic projection neurons in the basal forebrain in adulthood in male rats. In particular, we found increased WFA labeling within the OFC and medial PFC of AIE-exposed rats. We further determined that this increase in WFA labeling involves an increase in expression when surrounding PV interneurons, as a greater proportion (∼5%) of these neurons are surrounded by PNNs in AIE-exposed subjects relative to controls. As PNNs regulate PV excitability and plasticity (Chu et al., 2018), these findings provide a potential molecular mechanism by which AIE exposure can disrupt cortical synchrony and behavioral flexibility. Moreover, while AIE exposure did not alter expression of ChAT or PV interneuron numbers in either PFC or striatal regions, we found that ChAT-expressing projection neurons in the basal forebrain are reduced in adult males following AIE exposure as previously reported (Boutros et al., 2015; Ehlers et al., 2011; Swartzwelder et al., 2015; Vetreno et al., 2014). These findings highlight the region-specific effects of adolescent alcohol exposure that persist into adulthood.

### AIE exposure increases PNN density around PV interneurons in prefrontal regions

We found significant increases in WFA labeling in both the OFC and mPFC following AIE exposure. The increased labeling within the OFC was consistent with previous findings observed in mice (Coleman Jr et al., 2014), and we extended this finding to the mPFC (both PrL and IL) using both DAB-IHC and immunofluorescence. Moreover, we related this increase in PNNs to the PV interneurons they are known to ensheathe (Reichelt et al., 2019) via dual immunofluorescence. Interestingly, we found very few to no PNNs within the striatum or basal forebrain of our subjects regardless of group, suggesting that these nets are crucial for cortical, but not subcortical, neuronal regulation in adult rats. The lack of WFA labeling within striatum was unsurprising as several papers report low expression in rats (Bertolotto et al., 1996; Seeger et al., 1994), but not mice (Lee et al., 2012; see Sorg et al., 2016 for review).

As PNNs preferentially surround PV+ GABAergic interneurons in various cortical areas (Baker et al., 2017; Härtig & Brauer, 1992), we hypothesized that this increase in WFA labeling could be specific to this neuron population within the PFC. The present data indicate that in adult male rats, ∼30-40% of PV interneurons in the OFC and ∼10-15% of PV interneurons within the mPFC are surrounded by PNNs, and that the proportions of PV interneurons that were ensheathed by nets were significantly higher in AIE-exposed subjects in both the OFC and mPFC subregions. In contrast, we detected no co-localization between ChAT and WFA labeling within the PFC of sample tissue; therefore, no ChAT+/WFA+ co-label was conducted in AIE and CON tissue. This lack of WFA-ensheathed ChAT expression was expected as there are no reported instances of ChAT/PNN overlap in any CNS tissue (Brauer et al., 1993; Brückner et al., 1994; Morawski et al., 2014) although some report overlap within the spinal cord (Irvine & Kwok, 2018).

Together, our findings suggest that since PNNs that surround PV interneurons may influence the plasticity of these neurons (Chu et al., 2018; Reichelt et al., 2019), changes in their colocalization may contribute to suboptimal functional connectivity previously observed after AIE exposure (Broadwater et al., 2018; Gómez-A et al., 2021). Future studies should assess whether mechanisms that reduce PNN expression in the PFC (e.g., via breakdown by chondroitinase ABC (Brückner et al., 1998; Chu et al., 2018)) may ameliorate these deficits in functional connectivity and behavioral flexibility by restoring excitatory/inhibitory balance (Hensch, 2005).

### AIE exposure does not reduce numbers of prefrontal and striatal PV-expressing or ChAT-expressing interneurons

Of the various populations of neurons required for cognitive processes, cholinergic and GABAergic neurons may be particularly sensitive to the effects of adolescent alcohol exposure (Coleman Jr et al., 2011; Galaj et al., 2019; Vetreno et al., 2014). Changes in ChAT and PV interneuron populations or functions can have critical implications for AIE-induced disruption of behavioral output, as both populations are thought to coordinate and regulate medium spiny neuron organization during behavior (Gritton et al., 2019).

GABAergic PV interneurons constitute around 30-50% of the total number of inhibitory cells in the neocortex and participate in the control and regulation of cortical networks through local regulation and stabilization of excitatory activity (Batista-Brito et al., 2020). Thus, PV interneurons contribute to overall excitatory/inhibitory balance (Nahar et al., 2021) which impacts learning and memory behaviors (Murray et al., 2015). Moreover, these neurons regulate neural synchrony within microcircuits (Ruden et al., 2021) and, thus, potentially contribute to functional connectivity deficits observed by using resting-state MRI following AIE exposure (Broadwater et al., 2018; Gómez-A et al., 2021). Previous studies have demonstrated that AIE exposure in males reduces PV interneuron number in the hippocampus (Liu & Crews, 2017) but not basal forebrain (Coleman Jr et al., 2011). In the current study, we found no differences in PV-positive interneuron number in the mPFC, OFC, striatum, or basal forebrain; together with the previous reports, these findings suggest that PV neurons in the hippocampus maybe particularly vulnerable to AIE exposure effects (Liu & Crews, 2017).

Earlier studies reported that ChAT projection neurons in the basal forebrain (Vetreno et al., 2020) and ChAT interneurons in the dorsal striatum (Vetreno et al., 2014) and nucleus accumbens (Galaj et al., 2019) show reduced phenotypic expression in adult males after AIE exposure. While we replicated the loss of basal forebrain ChAT projection neurons, we did not observe any differences in populations of ChAT interneuron numbers in the striatum. To follow up on this discrepancy, we replicated the experiment in a different set of animals with similar results, suggesting that, under the conditions we used in the present study, ChAT interneurons in the striatum were not altered after AIE exposure. In addition, we found no loss of ChAT interneuron numbers in the mPFC and OFC after AIE exposure in males. One notable difference between our study and the Galaj et al. (2019) and Vetreno et al. (2014) studies was the inclusion of behavioral paradigms prior to tissue collection in the earlier studies, and it is possible that engagement of prefrontal circuits in behavioral training amplified effects of AIE exposure in those studies. Moreover, AIE-induced loss of ChAT projection but not interneuron numbers observed here could reflect greater sensitivity of projection neurons than interneurons to adolescent binge alcohol exposure. For example, maintenance of ChAT neuronal phenotype requires a complex interplay between transcription factors, proinflammatory factors, and trophic support, requiring activation of various receptors and intracellular cascades (as reviewed by Allaway & Machold, 2017). A critical difference between projection neurons and interneurons is that only projection neurons express the p75^NTR^ receptor, which exerts a pro-apoptotic effect. Therefore, ChAT projection neurons may be more sensitive to alcohol-related disruption of phenotype due to differences in p75^NTR^ activation (Vetreno & Crews, 2018). Importantly, while we saw no loss of ChAT expression in PFC interneurons, the significant loss of acetylcholine from basal forebrain projection neurons could disrupt PFC function and result in lasting behavioral inflexibility previously reported following AIE exposure (Galaj et al., 2019; Vetreno et al., 2020).

### Limitations and Future Directions

A notable limitation of the present study is that it included only male rats; thus, future studies of investigation should include females. This extension is particularly important as some studies show similar behavioral deficits in males and females while others show sex-specific deficits in behavioral tasks following AIE exposure (Macht et al., 2020; Madayag et al., 2017). Moreover, very little is known regarding sex differences in the development of PNN expression. However, it is possible that sex differences in the long-term neurochemical consequences of AIE exposure on PNNs will emerge, as expression of PNNs throughout adolescence develops in a sex-specific manner (Drzewiecki et al., 2020).

Another limitation of the present experiment is that only WFA labeling was used to mark extracellular components. As WFA is a general marker that binds to chondroitin sulfate N-acetylgalactosamine, it may have captured structures aside from PNNs. Future studies can assess more specific lecticans and link proteins, as many components of PNNs peak in expression at various developmental time points (Zimmermann & Dours-Zimmermann, 2008) resulting in lectican- and protein-specific AIE exposure effects (Coleman Jr et al., 2014). Finally, our histological analysis of WFA and PV was limited to expression of phenotypes rather than functional consequences; thus, future studies could determine if increased WFA labeling surrounding PV interneurons impacts the intrinsic excitability of those neurons and to provide insight into the functional relationship between PNNs and the neurons they ensheathe.

## Conclusions

The present data collectively suggest that the extracellular PNN component, WFA, is vulnerable to ethanol insult during adolescent development. Using co-labelled immunofluorescence, we determined that AIE exposure increases the percentage of PV neurons that were ensheathed by PNNs in the PFC. Overall, these findings highlight that adolescent alcohol exposure increases the aggregation of the PNN marker WFA around PV interneurons. This change to PNN integrity could potentially disrupt the intrinsic properties of the neurons they ensheathe resulting in altered frontostriatal circuitry related to behavioral deficits previously observed after AIE exposure.

