## Supplemental Figures for "Impact of adolescent intermittent ethanol exposure on interneurons and their surrounding perineuronal nets in adulthood"

**Supplemental Material**

*Authors:* Carol A. Dannenhoffer, PhD^1^ *, Alexander Gómez-A, PhD^1^ *, Victoria A. Macht, PhD^1^, Rayyanoor Jawad, BS^1^, E. Blake Sutherland^1^, Ryan P. Vetreno, PhD ^1,5^, Fulton T. Crews, PhD ^1,5,6^, Charlotte A. Boettiger, PhD ^1,2,3,4^, Donita L. Robinson, PhD ^1,4,5^

*co-first authors

Author affiliations:

^1^Bowles Center for Alcohol Studies, School of Medicine, University of North Carolina at Chapel Hill

^2^Department of Psychology and Neuroscience, University of North Carolina at Chapel Hill

^3^Biomedical Research Imaging Center, University of North Carolina at Chapel Hill

^4^Neuroscience Curriculum, University of North Carolina at Chapel Hill

^5^Department of Psychiatry, School of Medicine, University of North Carolina at Chapel Hill

^6^Department of Pharmacology, School of Medicine, University of North Carolina at Chapel Hill

Corresponding author:

Donita L. Robinson, PhD.

Bowles Center for Alcohol Studies, CB 7178

University of North Carolina at Chapel Hill, NC, 27599-7178, USA

**Supplemental Figure 1.**

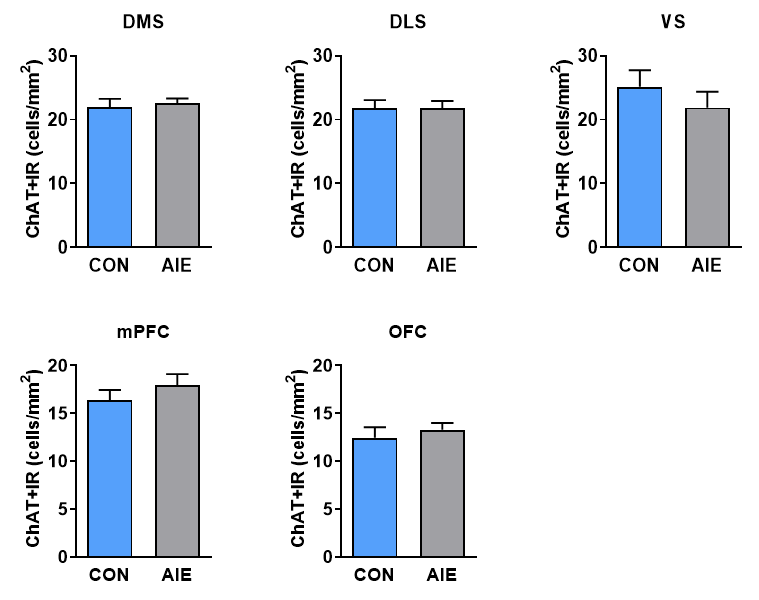

**Supplemental Figure 1. Adolescent intermittent ethanol (AIE) exposure did not alter striatal or prefrontal choline acetyltransferase (ChAT) expression.** T-tests were used to assess group differences within subregion; however, *α* levels were adjusted to correct for multiple comparisons (striatum: *α*=0.0167; PFC: *α*=0.025). Top panel: The striatum was subdivided into dorsomedial (DMS), dorsolateral (DLS), and ventral (VS) regions. ChAT immunoreactivity (ChAT+IR) was not significantly different between AIE and control (CON) subjects in the DMS (*p*=0.677), DLS (*p*=0.957), or VS (*p*=0.387). Bottom panel: The prefrontal cortex was subdivided into medial (mPFC) and orbitofrontal (OFC) regions. Neither of these subregions had differences in expression of ChAT (mPFC: *p*=0.335; OFC: *p*=0.515). Data are presented as mean ± SEM.

**Supplemental Figure 2.**

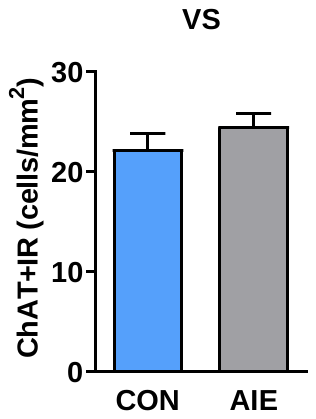

**Supplemental Figure 2.** Archived tissue from a previous study (CON n=8; AIE n=8) was used for follow-up analyses; that study used male Sprague-Dawley rats that underwent identical alcohol and water exposures. This follow-up analysis confirmed no change (*p*=0.255) in ventral striatal ChAT expression following AIE exposure. Data are presented as mean ± SEM.

**Supplemental Figure 3.**

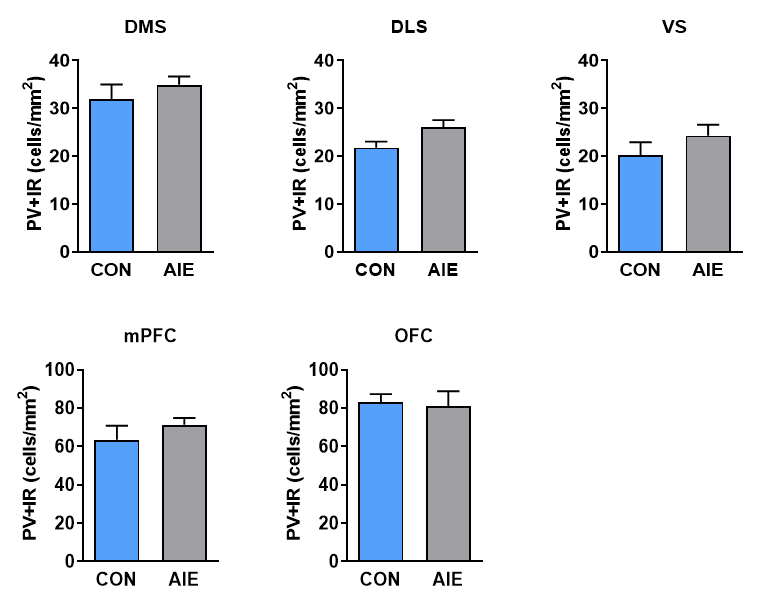

**Supplemental Figure 3. Parvalbumin (PV) expression is unaltered following AIE exposure.** T-tests were used to assess group differences within subregion; however, *α* levels were adjusted to correct for multiple comparisons (striatum: *α*=0.0167; PFC: *α*=0.025). Top panel: The striatum was subdivided into dorsomedial (DMS), dorsolateral (DLS), and ventral (VS). PV immunoreactivity (PV+IR) was not significantly different between AIE and CON subjects in the DMS (*p*=0.387), DLS (*p*=0.051), or VS (*p*=0.267). Bottom panel: The PFC was subdivided into mPFC and OFC; however, no difference in PV expression was detected in either subregion (mPFC: *p*=0.317; OFC: *p*=0.816). Data are presented as mean ± SEM.

**Supplemental Figure 4.**

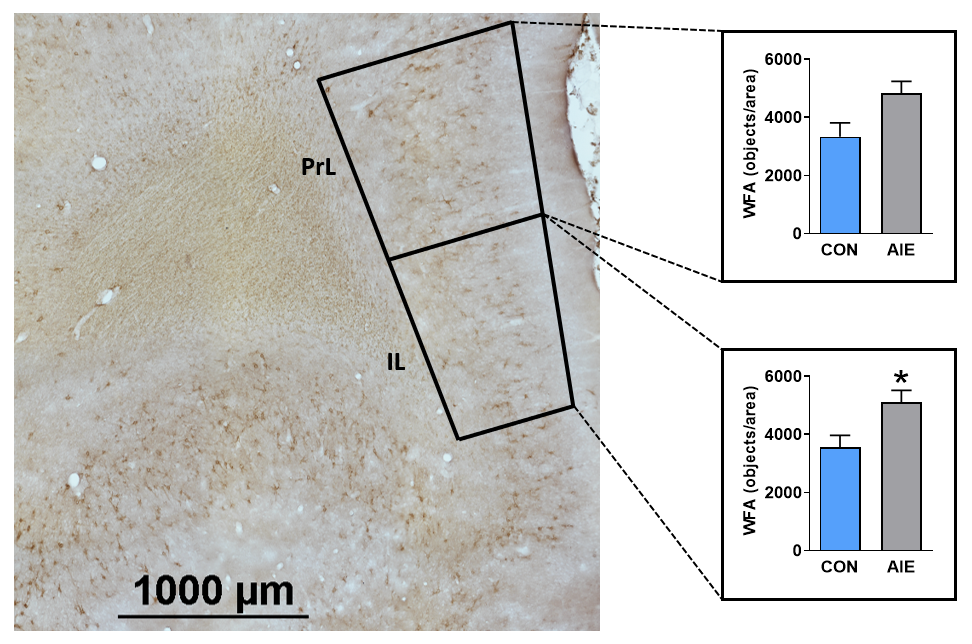

**Supplemental Figure 4.** We observed an increase in overall wisteria floribunda lectin (WFA) binding in the mPFC; hence we further subdivided this region into prelimbic (PrL) and infralimbic (IL) and adjusted *α* to correct for multiple comparisons (*α*=0.025). WFA expression was significantly higher within the IL of AIE rats (*t* (17) =2.63; *p*=0.02) and a trend was observed within the PrL (*p*=0.0255). Data are presented as mean ± SEM. * indicates main effect of exposure.

**Supplemental Figure 5.**

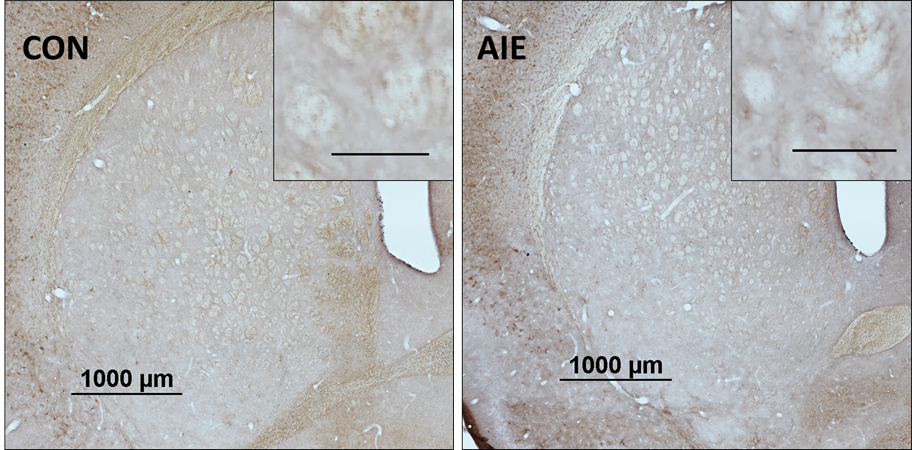

**Supplemental Figure 5.** We observed no WFA binding within the striatum of rats; therefore, data analysis was not conducted on this area. Scale bar in small inset image represents 100 µm.

**Supplemental Figure 6.**

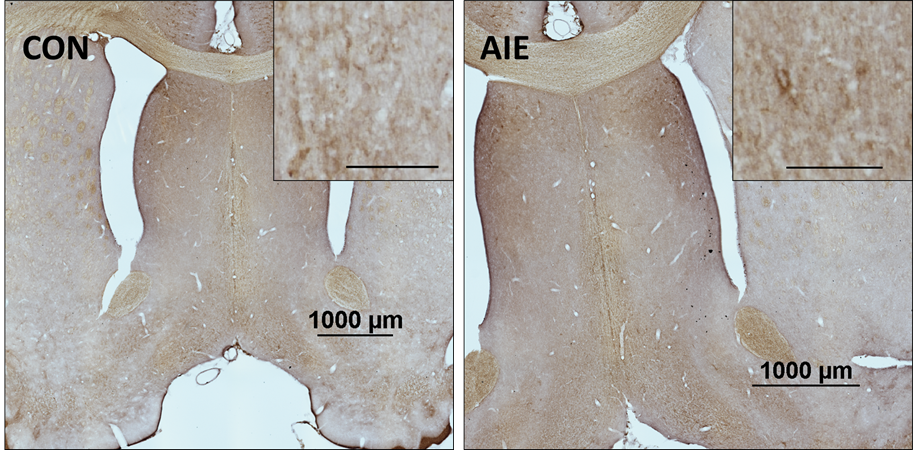

**Supplemental Figure 6.** We observed no WFA binding within the basal forebrain of rats; therefore, data analysis was not conducted on this area. Scale bar in small inset image represents 100 µm.
